# Computational alternative to cardiac biopsy in multiple myeloma: utilizing the shape of routine laboratory data for the classification of AL cardiac amyloidosis

**DOI:** 10.64898/2026.10.27.762298

**Authors:** Maegan A. Cremer, Hyeju Song, Michael G. Fradley, Isaac B. Rhea, Chris A. Kieslich, Meghan C. Ferrall-Fairbanks

## Abstract

**Background:** Patients with multiple myeloma (MM) are at increased risk for light chain cardiac amyloidosis (CA) due to high circulating levels of light chains. ALCA is a clinical diagnosis based on biopsy, cardiac imaging findings of restrictive cardiomyopathy, and serum monoclonal proteins. To overcome these challenges, we have developed machine learning models that accurately predict ALCA status while identifying key diagnostic features based on routine laboratory measurements and using limited data.

**Methods:** Utilizing electronic medical data (EMR) data, we built support vector machine models (SVMs) using the MISTIC framework. Data including diagnosis history, treatment, and laboratory values for patients with MM or monoclonal gammopathy of unknown significance (MGUS) were collected as a retrospective cohort study at Moffit Cancer Center and Research Institute between 2014 and 2019. To utilize the wealth of information in longitudinal EMR data, we fit regression models to the EMR data and used the parameters and characteristics of the fits as features for our SVMs. We tested three different sets of features: descriptive statistics alone, parameters and characteristics of fit only, and both the descriptive statistics and parameters of fit (DescFits). A classifier using raw data at a timepoint proximal to the date of ALCA diagnostic imaging (EchoAdj), as well as analogous random forest and XGBoost models were used for comparison.

**Findings:** SVM classifiers trained on the DescFits feature set (AUC= 1·0) outperformed blind model predictions of the EchoAdj model (AUC = 0·36) and other conventional machine learning approaches in identifying which patients were diagnosed with ALCA. The DescFits model selected features describing the temporal relationship of physiologic parameters including hematologic state, and cardiac and renal function.

**Interpretation:** We demonstrated that the MISTIC SVM framework and the shape of EMR data can aid in identifying patients with cardiac amyloidosis without cardiac biopsy or imaging.

**Funding:** This work was in part supported by funding from the National Institutes of Health (NCATS KL2TROO1429 to MCF and NIGMS R35GM147164 to CAK) and Moffitt Cancer Center and Research Institute’s Junior Scientist Partnership Award (to MCF and IBR). MAC is a trainee in the Medical Scientist Training Program supported by the National Institute of General Medical Sciences of the National Institutes of Health under award number T32GM163993.

## INTRODUCTION

Patients with multiple myeloma (MM) and other light-chain producing hematologic disorders (e.g. monoclonal gammopathy of unknown significance (MGUS)) are at increased risk for AL cardiac amyloidosis due to high levels of circulating light chains.^1,2^ Multiple myeloma is characterized by an abnormal monoclonal proliferation of plasma cells and increased circulating light chains.^3,4^ When these proteins misfold they can deposit throughout the body as amyloid light-chain contributing to organ dysfunction.^1,5–7^

Cardiac amyloidosis is the accumulation of amyloid in heart tissue.^8,9^ The extent of cardiac involvement, described as cardiac amyloidosis, is predictive of survival in light chain amyloidosis, where greater cardiac involvement is associated with reduced survival.^7,10–12^

AL Cardiac amyloidosis can present clinically as fatigue, conduction abnormalities and/or heart failure.^1,6,13–16^ Initial evaluation includes measures of cardiac function (blood pressure, NT-proBNP or BNP, troponin I or T), lytic bone activity (alkaline phosphatase), and kidney function (creatinine and 24-hour urine protein).^7^ The clinical diagnostic guidelines for cardiac light chain amyloidosis include serum positive for monoclonal proteins, amyloid positive biopsy at another site , and cardiac imaging findings of restrictive cardiomyopathy.^1,5,7,9,17^ Echocardiography measures can vary across technicians and instruments complicating assessment.^7^ If suspicion is high despite negative extracardiac biopsy, a higher risk endomyocardial biopsy is often required.^1,18719,20^ However, definitive diagnosis includes biopsy and testing for light chains to rule out other causes of cardiac amyloidosis.^6^ Treatment for AL cardiac amyloidosis is treating the underlying pathology.^1,7^ Patients with MGUS and smoldering myeloma may experience delays in treatment because of lower levels of circulating light chains and the difficulty of detecting AL amyloid presence. Therefore, for patients to receive timely care we need minimally invasive methods for determining patient risk for AL cardiac amyloidosis.

Due to the heterogeneity of clinical presentation, there is no unifying and singular set of definitive laboratory metrics or imaging measures for the determination of cardiac amyloidosis in patients with MM or MGUS. Our goal is to utilize machine learning (ML) models to determine a MM and MGUS patient’s cardiac amyloidosis status from routine laboratory measures without the use of radiographic measurements. Ultimately, this tool could be used as an additional tool to delineate patient risk for AL cardiac amyloidosis which serum and biopsy results are uncertain.

Classification algorithms have been developed as screening and diagnostic aids for a range of conditions including lung cancer, obstructive coronary artery disease, and cardiac amyloidosis in a general patient population.^18–21^ However, existing classification models for cardiac amyloidosis utilize cardiac imaging data or require use of imputation techniques for missing data.^18,19,22,23^ The missing not-at-random patterns often found in healthcare data are incompatible with most imputation techniques.^24–27^ For instance, a patient who is experiencing severe symptoms or rapid progression is more likely to have a greater frequency of contact with healthcare providers than a patient who is comparatively less limited by their health status.^24,27^

One major shortcoming of ML models for clinical applications is their black-box nature and lack of explainability.^28^ Recent efforts using large deep learning models have found improved predicted performance, but such models often compound challenges related to opaqueness of machine learned models and tend to require large amounts of training data. For conditions like ALCA in multiple myeloma patients, where clinical studies are often carried out with small patient cohorts, accumulating homogenous data at the scale needed to develop deep learning models can be challenging. Accordingly, there is a great need for data efficient machine learning frameworks to enable the development of accurate and explainable data-driven diagnostic models based on small clinical cohorts. In this paper we have applied a framework for building interpretable and explainable support vector machine models, MISTIC, to develop ALCA diagnostic models based on EMR data, including novel features that capture dynamic changes within in routine laboratory measurements. We also developed reference models, specifically random forest and XGBoost models trained on the full feature set or using data only near ALCA diagnosis, to represent standard approaches in literature. We report the performance of all of these models, evaluate the usefulness the feature types and multiple feature combinations, and describe the potential for clinical translation of our models based on the identified diagnostic features.

## METHODS

### 1. Ethics statement and data acquisition

Participants with multiple myeloma (MM) or monoclonal gammopathy of unknown significance (MGUS) who were evaluated for cardiac amyloidosis (CA) were recruited at Moffitt Cancer Center and Research Institute between December 2014 and January 2019 under IRB Protocol MCC19770 for this retrospective cohort study. The temporal window of data is 60 days before oncologic diagnosis until completion of the data collection window. An initial cohort of 48 patients with 24 ALCA+ and 24 ALCA-patients were identified. Data included demographic information, diagnosis, treatments received, physician note information (clinical impression of disease, ECOG scores, Karnofsky scores), laboratory information contained in the medical records, and cardiac imaging data with interpretations by a cardio-oncologist. All data were de-identified and time indexed to the oncologic date of diagnosis. Patients were excluded from the dataset for missing greater than 80% of labs (data length 0) and/or data length 0 for kappa-free light chains, troponin-T, or NT-proBNP. This resulted in a final cohort size of 44 patients, 21 ALCA+ and 23 ALCA-, with demographic information reported in **Figure 1A** and an event plot describing incidences of laboratory data across time in **Figure 1B**. Distribution of key echocardiographic measures, distribution of laboratory data proximal to the date of diagnostic imaging, heat map of number of incidences of each lab for each patient, and distribution of average lab values are reported in **Supplemental Figures S1-S4.**

**Figure 1.**
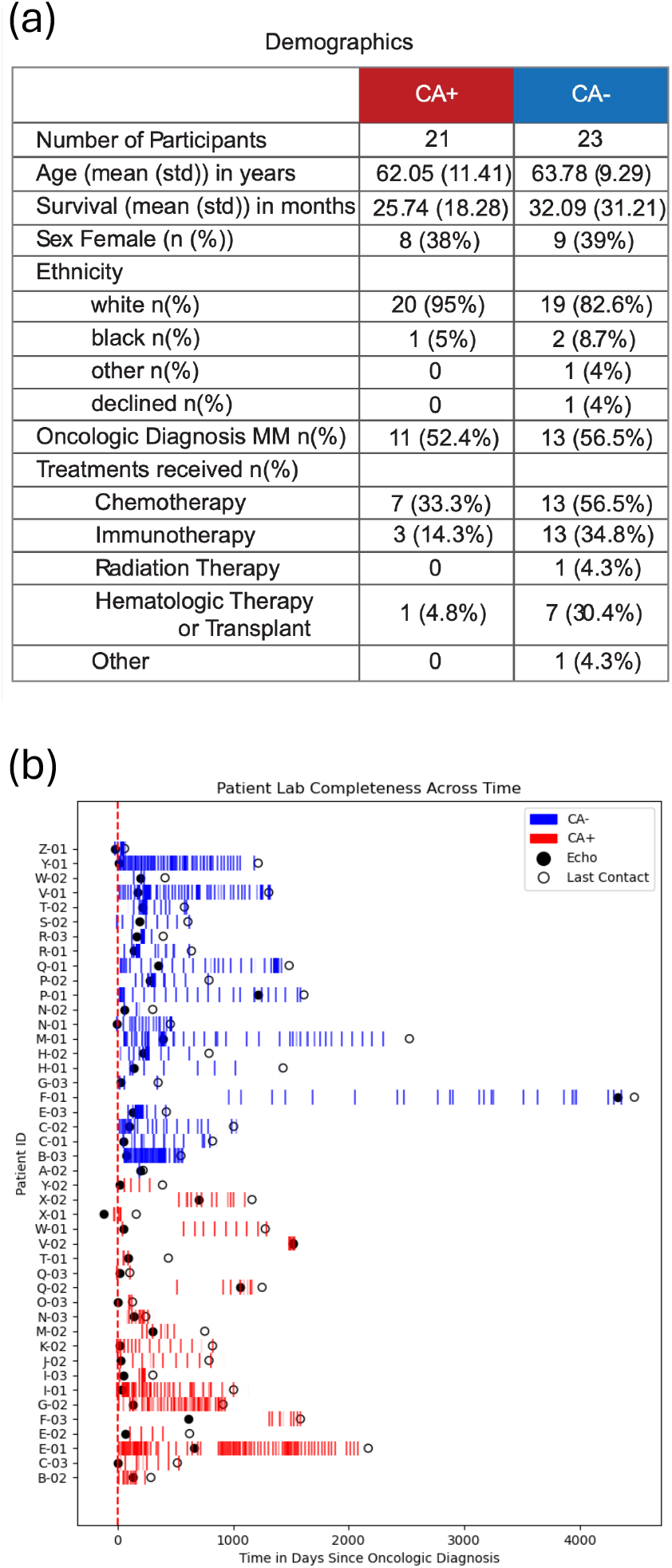
Cohort Demographics and Data Completeness. Patients were recruited between 2015-2019 at Moffitt Cancer Center and Research Institute who were diagnosed with multiple myeloma (MM) or monoclonal gammopathy of unknown significance (MGUS) who were evaluated for cardiac amyloidosis (CA). Patient demographics are listed in table (A) including oncologic diagnosis and treatments received. The completeness of patient data is shown in the event plot in panel (B) where the opacity corresponds to the completeness of the 39 common labs across all patients.

### 2. Data pre-processing and encoding Selection of laboratory measures

Data was available for 59 routine laboratory (lab) values collected across all patients. The list of labs used in our models was reduced to the 39 labs that were captured for all patients with a data of length 1 or greater listed in **Supplemental Table S3**. This includes metabolic panel results (e.g., creatinine, BUN, sodium, chloride, calcium), complete blood count with differential (e.g., WBC and RBC counts, MCV, hematocrit), circulating protein levels (e.g., albumin, kappa free light chains), serum electrophoresis, inflammatory markers (e.g., LDH, ferritin), and cardiac markers (e.g., NT-proBNP and troponin-T). The distribution of lab data density across time are described by the event plot in **Figure 1B**, with the number of recorded instances of each lab and the average value of a lab by amyloid status in **Supplemental Figure S3 and S4**.

#### Encoding of Categorical Variables and Treatment Information

Categorical data including cardiac amyloidosis status, sex, and cancer diagnosis (multiple myeloma, MGUS, or other) were encoded using the one-hot encoding style of “get_dummies()” function in Python Pandas.^29^ This created individual columns or binary features for each category. The column for cardiac amyloidosis positive (where positive is 1) was used for the outcome variable in the classifier. Columns containing duplicate binary information were dropped from the dataset including the cardiac amyloidosis status negative column (negative = 1), male sex (male = 1), and diagnosis other (other = 1).

#### Encoding Physician Note Information and Determining Clinically Relevant Test-Train Split

Clinical scoring metrics for a patient’s functional status include Karnofsky Performance Scores (KPS) and ECOG scoring.^30^ An equivalency was established between the KPS and ECOG systems as described in **Supplemental Figure S5**. Because ECOG scoring is 0-5 (lower is more independent) in increments of 1 while KPS scoring is 100-0 (higher is more independent) in increments of 10, converting from ECOG to KPS scoring establishes an ECOG score to an intermediate KPS (ECOG 0 is KPS 95). A KPS score is mapped to the nearest ECOG score (KPS 80 and KPS 70 are ECOG 1). After the data was completed using the ECOG to KPS equivalencies, a mean score was calculated per patient across the relevant time window. A 1/5 test-train split was created using the distribution of mean Karnofsky scores across cardiac amyloidosis positive and negative patients. The first and every fifth patient was selected from the ALCA+ and ALCA-distributions to form the test set. One ALCA+ patient was missing both Karnofsky and ECOG scores and was assigned to the test set. The distribution of mean KPS scores for all patients and the train-test groups are plotted in the **Supplemental Figure S5**.

### 3. Dynamic Feature Generation from Lab Data

To honor the temporal relationships of patient lab data, time coarse data for each patient lab type were fit to model equations (linear and exponential) and their parameters (coefficients, intercepts, fit quality) were included as patient features. This enables all features of a patient’s data across time to be handled as a single instance by the model and reduced the missingness of data without using imputation. Additionally, classical statistical descriptions of data distributions (mean, median, quartile ranges, standard deviation) were included as features of a patient’s data. Participant lab data for each of the 39 labs, were fit to linear and exponential equations using the “LinearRegression” package from SciKitLearn in Python^31^ For fitting to the exponential equation *y* = *A* ∗ *e^Bx^*, the data was first translated by 1 to account for true zeros in cell count measures, then transformed using the natural logarithm and fit to a linear equation *ln*(*y*) = *ln*(*A*) + *Bx*. Data of length 2 were required for linear and exponential models (after transformation using the natural logarithm). The quality of fit was reported as R^2^ (coefficient of determination), residual error, and descriptive statistics for the residuals between the model and original data. Data of length 1 were resolved across all models by setting the y-intercept to the recorded value, the slope(s) to 0, and the R^2^ to -1 describing a “poor or under-informed fit,” since the true trajectory could not be known. For data of length 1, the descriptive statistics were set to the participant’s single recorded value. Features corresponding to linear fits were labeled linear, while features corresponding to exponential fits were labeled exp. Example plots of patient data and corresponding regression models are shown in **Supplemental Figure S6.**

### 4. Model Conditions/Data and MISTIC SVM construction

The MISTIC framework leverages support vector machines (SVM) due to their suitability for small datasets and their inherent interpretability as they identify samples (patients) that represent the boundary between the two classes (diagnoses)^32,33^. MISTIC builds on top of the support vector classifier (SVC) provided by SciKitLearn and provides novel tools for feature selection that account for how features contribute to separation of the two classes (diagnoses) at the boundary, as well, how features contribute to the overall spread of the samples. MISTIC also provides interpretations in the form of feature ranking, and explanations based on integrated gradients, which measure how much a patients prediction score is due to specific feature values.

We applied the MISTIC framework to train SVM models using different initial feature sets: (1) EchoAdj model using a slice of raw patient data proximal to time of diagnostic cardiac imaging (median 29.7 days and 59.9 days prior to imaging among ALCA+ and ALCA-patients respectively, the relative time for imaging and adjacent labs are listed in **Supplemental Table S2**), (2) DescOnly model using only the descriptive statistics of patient data, (3) ParamsOnly model using only the parameters and characteristics of fit of patient data, and (4) the DescFits model using both descriptive statistics and parameters of fit as model features as shown in **Figure 2**. All models included demographic and treatment information. Missing data in the EchoAdj model was assigned a value of -1 because 0 has physiologic importance.

**Figure 2.**
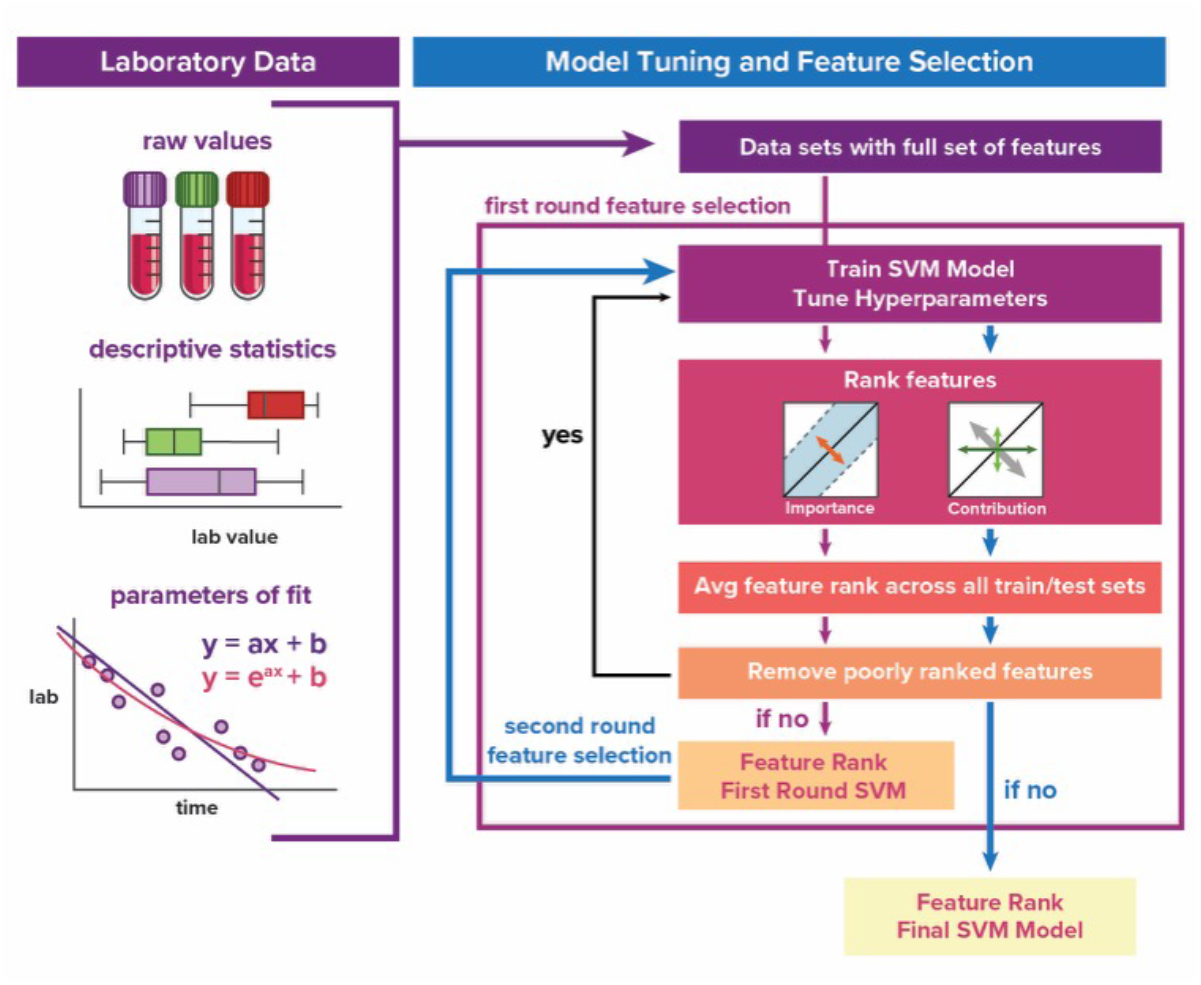
Methods Illustration of Data Processing and Feature Selection using MISTIC SVM. The initial data includes demographic and treatment information as well as routine EMR data. This data presented to the model as features in three ways: raw values at a single point in time per patient, descriptive statistics over time, and/or parameters of fit to model equations (linear and exponential). Using backwards feature selection and the MISTIC framework for feature ranking, features are removed from the model until there are no poorly ranked features. The final model has the greatest performance with the fewest number of features. This process is repeated a second time to refine the model’s feature list.

Data was transformed using the normal scalar prior to training models. A group of five ALCA+ and five ALCA-cases were withheld as a training set as described in “ **Encoding Physician Note Information and Determining Clinically Relevant Test-Train Split”**. The remaining cases were used for 5-fold cross-validation. Hyperparameter tuning and feature selection was conducted using MISTIC based on a precomputed linear kernel with cost values (C range -16 to 10)^33^. Briefly, a backward feature selection process was conducted across two rounds with feature ranking using weights of importance and contribution as described in **Figure 2**. The best performing model among the different weights of importance and contribution was determined using an enrichment score, which measured the average performance across the feature selection process. In this work, the average of area under the receiver operator curve (AUC ROC) and F1 score were used to measure performance during hyperparameter tuning and feature selection. Here we combined both scores, since F1 score measures the model’s ability to correctly place the patients in their diagnostic category and AUC ROC measures the model’s ability to sort patients into categories. In the first round of feature selection, 25% of remaining features were removed after each iteration and successive iterations of feature removal were applied until only a single feature remained. Based on the best performing feature set from round one, a second round of feature selection was applied in which a single feature was removed in each iteration, and the best performing model and features from round two represent our final models. After model training and feature selection using MISTIC, the resulting models were evaluated on the blind prediction sets using ROC plots.

### 5. Comparison of MISTIC SVMs to Other Classification Methods

To compare the MISTIC SVMs to conventional machine learning classifiers, random forest and XGBoost models utilizing feature selected by the analogous SVM models were constructed. To test the contribution of the random forest architecture vs the feature list in model performance, a random forest model was created using its top ranked features of length equivalent to the corresponding SVM. The random forest and XGBoost models were evaluated using the identical test set of patients used for the MISTIC models and performance was measured using AUC ROC.

### 6. Statistical Analysis

The statistical tests were conducted in GraphPad Prism (v10·0·2) and Python (3·12·4 with packages: matplotlib 3·9·1^34^ , numpy 1·26·4^35^ , openpyxl 3·1·4, pandas 2·2·2^29^ , plotly 5·22·0, scikit-learn 1·5·1^31^, scipy 1·14·0^36^, seaborn 0·13·2^37^, shap 0·45·1^38,39^, xgboost 2·1·4^40^).^29,31,34–40^ To analyze statistical differences between groups in EchoAdj slice of raw lab values we used the non-parametric Mann-Whitney U test. To analyze the statistical differences between groups in the mean of lab values across time we used the multiple non-paired t-test with Holm-Sidak. To analyze statistical differences between groups in the mean Karnofsky scores we used the two sample KS test. For examination of the statistical correlation between selected features, we used the Spearmans correlation. DeLong tests for comparing the AUCs of machine learning models were ran in Python 3·13·11 using the MLstatkit (version v0·1·91).

### 7. Data and Code Availability

The code, data, and documentation of the machine learning models explored in this analysis are open-source and publicly available through a virtual machine producing the full computational environment available in a Code Ocean Capsule at: https://codeocean.com/capsule/6012685/tree. This repository includes a copy of the specific version of the MISTIC framework used in this work as the GitHub repository for MISTIC is under continuous development.

## RESULTS

### The simplified DescFits MISTIC SVM was the best performing model among the SVMs

Four different MISTIC SVM classifiers were constructed as shown by the purple path in **Figure 2**: (1) using only data adjacent to cardiac amyloidosis diagnostic imaging (EchoAdj), (2) only descriptive statistics of longitudinal data (DescOnly), (3) only parameters and characteristics of fit of longitudinal data (ParamsOnly), and (4) both descriptive statistics and parameters and characteristics of fits (DescFits). After the first cycle of feature selection (feature reduction rate of 25%) and hyperparameter tuning, all models had strong cross validation performance shown in **Figure 3A**. When comparing the number of features and support vectors in the first-round models (**Figure 3B**), the ParamsOnly 8 feature model had the lowest number of features (8), but the highest mean number of support vectors (20·4), indicating that approximately 2/3 of the training set was used to construct the hyperplane. The EchoAdj 21 feature and DescFits 23 feature models had similar performance to the ParamsOnly 8 feature model with AUC of 1·0 but have lower mean number of support vectors (16·8) and greater number of features. To better identify which features and support vectors were necessary for the final MISTIC SVMs, a second round of feature selection examining each model’s selected features was performed as described by the blue path in **Figure 2**.

**Figure 3.**
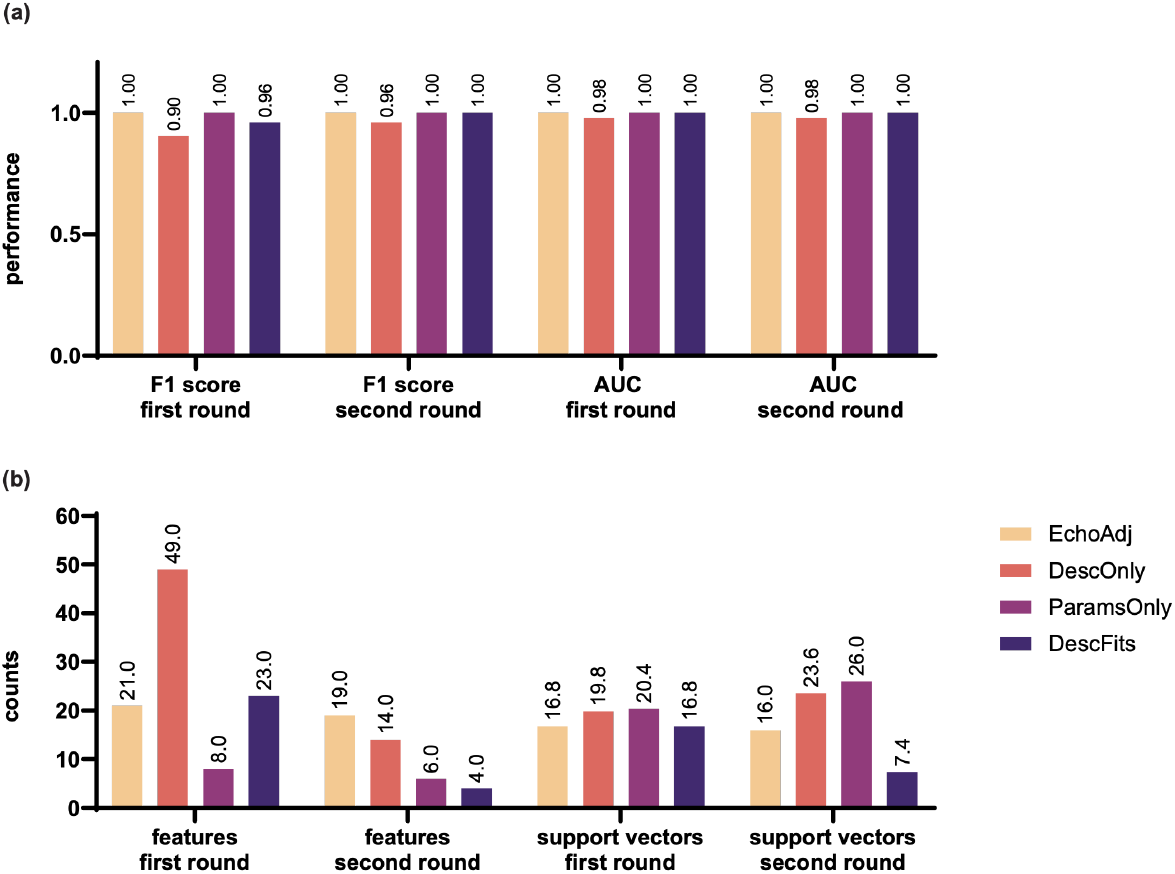
Cross-validation performance of first and second round SVM models. The cross-validation performance (F1 and AUC-ROC) of the first and second round MISTIC SVM models are reported in panel a. The number of features and support vectors for the first and second round MISTIC SVMs are reported in panel (b). The DescFits second round MISTIC SVM has the best-cross validation performance, lowest number of features, and fewest support vectors.

After the second round of feature selection (feature reduction 0), the cross-validation performance of the MISTIC SVMs was maintained as shown in **Figure 3A**. These simplified models were then tested on the blind set of 10 patients with ROC curves shown in **Figure 4**. All simplified MISTIC SVMs utilizing the shape of longitudinal data—either as traditional descriptive statistics and/or parameters of fit—outperformed data from a single point in time (EchoAdj 19 feature model blind AUC 0·36) when challenged with the blind test set.

**Figure 4.**
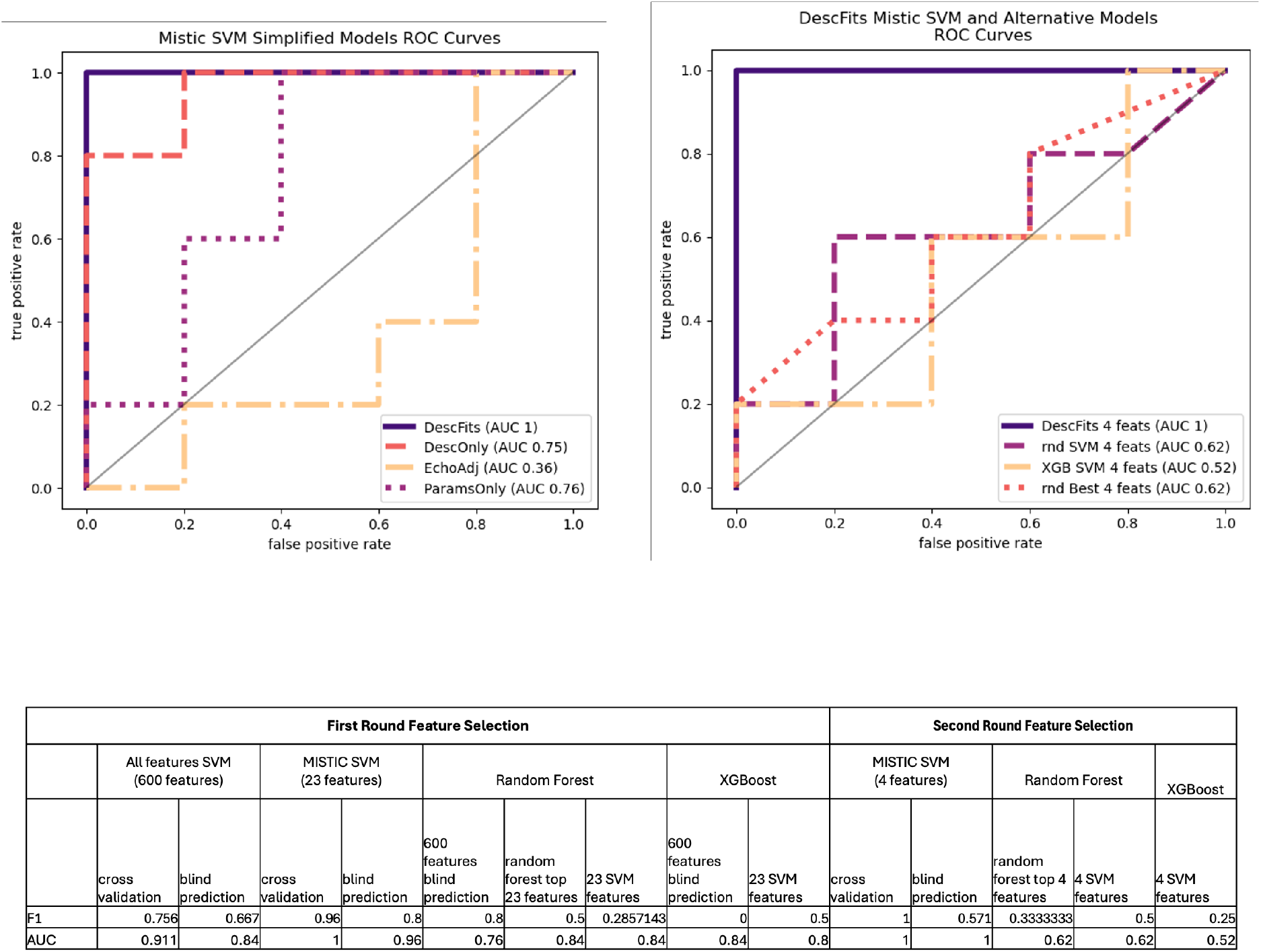
Model performance on the blind set. Receiver operating curve (ROC) using the blind test set for the second round MISTIC SVMs (top left) and the corresponding DescFits alternative models (top right). The cross-validation and blind prediction performance (F1 and ROC-AUC) of models using the initial 600 DescFits features, 23 MISTIC SVM selected features from the first round, and 4 MISTIC SVM selected features from the second round are reported in the table (bottom). Abbreviations: rnd svm, random forest model using 4 features selected by the DescFits SVM; rnd best, random forest using its top 4 features from the 23 feature SVM model; xgb svm, XGBoost using the 4 features selected by the DescFits 4 feature model.

The EchoAdj 19 feature model used a mean of 16 support vectors indicating that a high percentage of the training data was necessary to form the decision boundary. This makes the EchoAdj 19 feature model—and other models with a high ratio of support vectors to training data—less accommodating to noise in the data.

The DescFits 4 feature model successfully classified the test patients as indicated by the AUC of 1 (95% CI 1·0-1·0) **Figure 3 and 4**, out-performing the other MISTIC SVM models on the test set. When comparing the final number of features and mean number of support vectors seen in **Figure 3B**, the DescFits 4 feature model approximated the desired 10:1 ratio of samples to selected features. The lower number of support vectors used by the DescFits 4 feature model (**Figure 3B**) reduces the risk of over fitting to the training data.

### The simplified DescFits model outperformed the similarly constructed XGBoost and Random Forest models·

To evaluate the impact of the MISTIC framework on model performance, XGBoost and random forest classifiers were created using the features selected by the DescFits 4 feature model. A random forest classifier trained on its top 4 features ranked from the initial 23 features of the first DescFits model was created to evaluate the impact of the feature selection process on final model performance. The performance of the alternative models on the blind test set is shown as an ROC plot in **Figure 4B** and their feature rankings are listed in **Supplemental Table S3**. The DescFits 4 feature model (AUC 1, 95% CI 1·0-1·0) outperformed the XGBoost (AUC 0·52, 95% CI 0·105-0·935) and random forest (AUC 0·62, 95% CI 0·224-1·0) using the same set of features as well as the random forest on its best 4 features (AUC 0·62, 95% CI 0·239-1·0). The AUCs are statistically significantly different between the DescFits 4 feature model and XGBoost with SVM 4 features (z-score -2·268, p-value 0·023) and statistically insignificant for other comparisons due to the width of the random forest AUC confidence intervals. The best 4 features according to the random forest model included BUN and chloride features, two of these features were shared with the DescFits 4 feature SVM model. It is therefore likely that the performance of the DescFits 4 feature MISTIC SVM model can be attributed both to model architecture and selected features·

### The simplified DescFits MISTIC SVM selected 4 features describing cardiac function, renal function, and hematologic state·

In the second round of feature selection, the backward selection process evaluated each of the features selected in the first round of feature selection ranking their importance to the model decision. The feature ranking and progressive cross validation performance of the initial 23 features selected by the DescFits 23 feature model is shown in **Table 1**. There was an abrupt decrease in the mean number of support vectors between the 6 and 5 feature models and a slight increase in performance indicating that these 5 features are among the most descriptive of the dataset. There is no change in performance between the 5 and 4 feature models, thus the 4-feature model was chosen as the final DescFits model.

**Table 1.** DescFits feature ranking and relative model performance. Features are ranked from highest to lowest (top to bottom) from the 23 features selected in the first round of feature selection. Each subsequent model includes the features listed above it. The 4-feature model was found to maximize performance, while minimizing the number of features and support vectors.

| Feature | F1 | ROC-AUC | C value | Numb. Of features | mean numb. support vectors |
| --- | --- | --- | --- | --- | --- |
| Chloride linear intercept | 0.653 | 0.711 | 1.000 | 1 | 19.8 |
| BUN 50% | 0.900 | 0.956 | 16.000 | 2 | 10.4 |
| NT Pro BNP exp intercept | 0.900 | 0.956 | 16.000 | 3 | 9.8 |
| Serum Electrophoresis Beta linear intercept | 1.000 | 1.000 | 4.000 | 4 | 7.4 |
| Pct Immature Gran Auto exp mse | 1.000 | 1.000 | 16.000 | 5 | 5 |
| Beta-2-Microglobulin 75% | 0.960 | 0.956 | 0.016 | 6 | 27.8 |
| BUN linear r2 | 0.960 | 0.956 | 0.016 | 7 | 27.2 |
| Chloride mean | 0.960 | 0.956 | 0.016 | 8 | 25.4 |
| Platelet Count exp coef | 0.960 | 0.978 | 0.016 | 9 | 25.4 |
| WBC exp coef | 0.960 | 0.978 | 0.016 | 10 | 25.8 |
| total CO2 50% | 0.960 | 0.978 | 0.031 | 11 | 21.4 |
| Platelet Count linear coef | 0.960 | 0.978 | 0.016 | 12 | 25.4 |
| Nucleated RBC min | 0.960 | 0.978 | 0.031 | 13 | 20.8 |
| total CO2 mean | 0.920 | 0.978 | 0.031 | 14 | 20.6 |
| Serum Electrophoresis Alpha 2 exp coef | 0.960 | 0.978 | 0.063 | 15 | 17.8 |
| total CO2 25% | 0.960 | 0.978 | 0.063 | 16 | 18 |
| Chloride 25% | 0.960 | 0.978 | 0.125 | 17 | 13.6 |
| Serum Electrophoresis Beta max | 0.960 | 0.978 | 0.125 | 18 | 13.8 |
| Potassium exp r2 | 0.960 | 0.978 | 0.125 | 19 | 14 |
| Pct Mono exp coef | 0.960 | 0.978 | 0.063 | 20 | 16.6 |
| Lambda Free Light Chains linear mse | 0.960 | 1.000 | 0.063 | 21 | 16.8 |
| Lambda Free Light Chains 50% | 0.960 | 1.000 | 0.063 | 22 | 17.2 |
| Serum Electrophoresis Alpha 2 linear intercept | 0.960 | 1.000 | 0.063 | 23 | 16.8 |

The initial 23 features from the first round of feature selection DescFits model, listed in **Table 1**, after the first round of feature selection included descriptions of cardiac function (NT pro BNP), renal function (chloride, potassium, BUN), anemia (nucleated RBCs), and hematologic state (serum electrophoresis beta, lambda light chains, WBC, % monocytes, platelet counts). As part of the feature narrowing, highly correlated features such as the platelet count exponential and linear coefficients (**Supplemental Figure S7**) were removed from the model. Anti-correlated features (correlations shown in **Supplemental Figure S7**) such as the exponential coefficient and linear intercept for serum electrophoresis alpha 2 were also removed from the model.

The final 4 features of the DescFits 4 feature model captured cardiac stress (NT pro BNP), renal function (BUN and chloride), and hematologic status (serum electrophoresis beta). To visualize how these 4 features were used to classify patients, the feature values were plotted against the feature’s integrated gradient with the point color determined by a second feature’s value as shown in **Figure 5** and **Supplemental Figure S8.** Positive integrated gradient values contributed towards a positive decision value classifying a patient as ALCA+.

**Figure 5.**
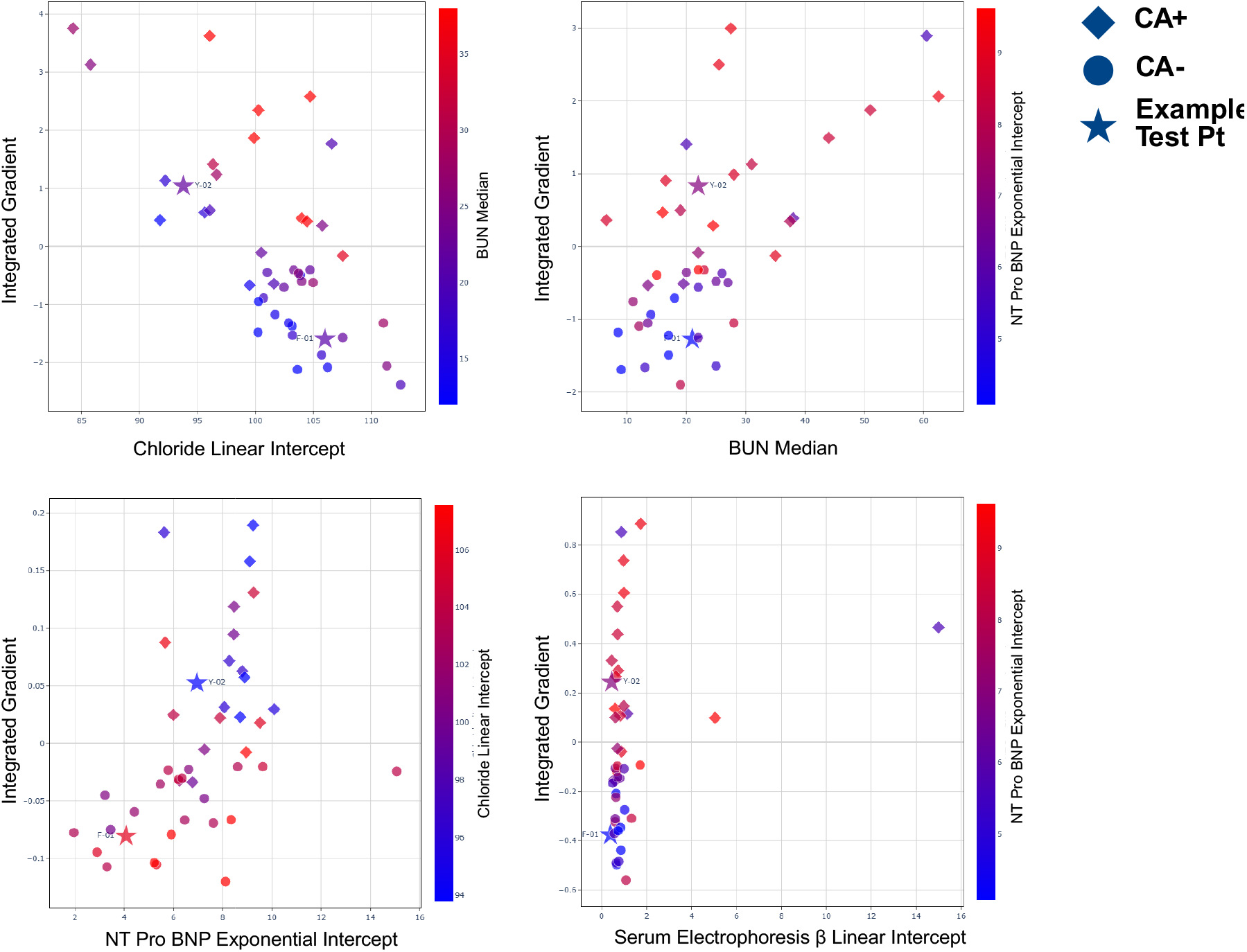
DescFits selected features integrated gradient plots. Integrated gradients vs feature values for each of the 4 selected features by the DescFits model: chloride linear intercept, BUN median, NT Pro BNP exponential intercept, and serum electrophoresis β linear intercept. The color of the plots reflects a secondary feature value as indicated. All patients are shown on the graphs. ALCA+ patients are marked with a diamond, ALCA-patients with a circle, and two representative test patients are marked with stars. Patient Y-02 is ALCA+ and patient F-01 is ALCA-. Additional combinations of secondary features are shown in supplementary figure 8.

Higher chloride linear intercept values (within the physiologically normal range for this set of patients) and lower BUN median values were associated with lower integrated gradient values and thus a ALCA-status as seen in **Figure 5**. This could describe patients with relatively well functioning kidneys. The separation between ALCA+ and ALCA-patients was further clarified by the integrated gradient plot of NT Pro BNP exponential intercept, a measure of cardiac stress. Higher chloride linear intercept values were associated with lower NT Pro BNP exponential intercept and negative integrated gradient values (for the BNP feature). This could describe a relationship of declining renal function and increased cardiac stress in our ALCA+ cohort consistent with the pathophysiology of cardiac amyloidosis.^6,7^

The beta region of the serum protein electrophoresis is sometimes discussed as a location for the M-spike in patients with multiple myeloma.^41–43^ The value of serum electrophoresis beta linear intercept alone was not significantly different by ALCA status. However, when examining the integrated gradient of this feature and overlaying the NT Pro BNP exponential intercept, we can clearly see a strong separation of the ALCA+ and ALCA-patients. This illustrates the complex relationship between the features that are captured by the SVM architecture.

### The function of model intercepts (projection to time 0) over raw values adjacent to the time of oncologic diagnosis or cardiac imaging

When fitting patient data to model equations (linear and exponential), we projected the intercept value to time 0, the date of oncologic diagnosis. The projected intercept value is influenced by data resolution. For example, patients with data far from time 0 (date of oncologic diagnosis) or a long-time course of data frequently have y-intercepts that are influenced by the shape of patient data over time (example plots of the test patients shown in supplemental figure 6). Additionally, the proximity of the date of diagnostic imaging to the date of oncologic diagnosis varies across patients and ALCA status (supplemental table S2). To evaluate the role of intercept features in the performance of the DescFits 4 feature model, we substituted the intercept features with the value nearest time 0 (DescFits-Onco) or the value nearest diagnostic imaging (DescFits-Echo). The cross-validation performance of the DescFits-Onco (AUC 0·867, F1 0·813) and DescFits-Echo (AUC 0·822, F1 0·786) models are weaker than the DescFits 4 feature model (AUC 1·0, F1 1·0). The blind performance of the DescFits 4 feature model (AUC 1·0, 95% CI 1·0, 1·0) out-performs the DescFits-Onco (AUC 0·72, 95% CI 0·408-1·0) and DescFits-Echo (AUC 0·68, 95% CI 0·265-1·0) models. The DeLong tests for statistically significant differences in AUC between machine learning models showed no statistically significant difference between the AUCs of the DescFits 4 feature models and DescFits-Onco (z -1, p-value 0·317) or DescFits-Echo (z -1·512, p-value 0·131) models due to the wide confidence intervals of the DescFits-Onco (AUC 95% CI 0·408, 1·0) and DescFits-Echo (AUC 95% CI 0·265, 1·0) models. This supports the assertion that the model intercepts are important for capturing the heterogeneity of patient temporal data.

## DISCUSSION / CONCLUSIONS

Patients with MM have an increased risk for AL cardiac amyloidosis (ALCA) due to high circulating levels of amyloid light chain. ALCA can present with a range of symptoms including fatigue, shortness of breath, limb swelling, and electrical abnormalities.^1,6,19,44^ While MM has a 5-year survival of about 60%, the progressive heart failure from light chain ALCA has a 20% mortality rate within 6 months.^5^ Due to the body’s inability to clear amyloid light chain, the effects of ALCA persist beyond multiple myeloma survivorship. Early detection and treatment for ALCA targeting the underlying plasma cell dyscrasia is essential for survival. However, the overt kidney dysfunction is often the first clinical entry to screening for cardiac amyloidosis due to the frequency of biopsy methods for diagnosing renal pathology.^6,7^ Cardiac biopsy is currently the gold standard diagnostic measure for ALCA.^5,6,19,23,44,45^ We aimed to determine a patient’s ALCA status using routine laboratory data without the need for cardiac biopsy or imaging data.

We created SVM models to classify patients with MM by their ALCA status using features derived from their EMR data. These SVM models capture the longitudinal shape of the data through the parameters and characteristics of fit to exponential and linear model equations. We found that the SVM model including both traditional statistics and the parameters and characteristics of fit (DescFits 4 feature model) was the best performing model (Cross validation AUC =1, blind prediction AUC = 1) across all SVMs tested (**Figure 3**) as well as analogous random forest and XGBoost models (**Figure 4**). This model accurately classified our patient set into ALCA+ and ALCA-groups without utilizing imaging ^22^, ECG ^20^ , or proteomic^46^ data commonly used by other models in the literature or clinical practice.^20,22,46^ The features selected by the DescFits 4 feature model (**Table 1**) captured elements of renal function, cardiac stress, and hematologic status. Three out of 4 of the features included parameters and characteristics of exponential or linear fits, indicating that the shape of EMR data was important for classification in this model. While the shape features are intercept values, the intercepts are influenced by the shape of the data over time and are not functionally equivalent to the measure nearest time 0 (DescFits-Onco) or diagnostic imaging (DescFits-Echo) for model performance. Our SVM DescFits model was unique because it utilizes features describing the shape of routine laboratory data collected as part of a patient’s oncologic care, does not require advanced tests or imaging (echocardiogram, cardiac MRI, ECG, or proteomic data), is computationally light, and captures clinical features relevant to the pathophysiology of cardiac amyloidosis progression. With further evaluation on larger datasets, this method has the potential to reduce the need for cardiac biopsy in the diagnostic pathway for ALCA in MM patients.

Utilizing EMR data collected from 44 patients undergoing evaluation for cardiac amyloidosis, we developed SVM models with linear kernels utilizing weightings of importance and contribution for attribution using the previously established MISTIC algorithm.^32,33^ To capture the shape and longitudinal relationships of EMR data, we fit patient data to exponential and linear model equations. The parameters and characteristics of fit were given as features to the SVM in addition to traditional descriptive statistics. The performance of the DescFits model was compared to SVM models trained on a section of raw data proximal to diagnostic imaging (EchoAdj), descriptive statistics alone (DescOnly), and parameters and characteristics of fit only (ParamsOnly). Additional comparisons were made to random forest and XGBoost models trained on the full feature list and the selected best features from each of the four aforementioned SVM models. Final versions of the MISTIC SVM models were created using a second round of feature selection from the initially selected features. Model performance was evaluated using F1 and AUC scores during cross validation reported in **Figure 3** and AUC ROC plots for blind prediction performance seen in **Figure 4**.

The DescFits 4 feature model successfully classified patients ALCA status (cross-validation F1 = 1, AUC = 1; blind prediction performance AUC =1) selecting the fewest features and utilizing the fewest support vectors as described in **Figures 3 and 4**. While an F1 and an AUC of 1·0 would raise concerns for overfitting, the DescFits 4 feature model maintains performance on the blind set (AUC =1, **Figure 3**) suggesting that this model will perform well on future datasets. The other SVM models displayed a decreased performance on the blind prediction set (**Figure 3**), consistent with over-fitting. This remarkable performance was greater than seen in competing models in the literature using imaging values.^18,22,23^

The DescFits 4 feature model also outperformed the analogous random forest and XGBoost models using the same list of features or a list of equivalent length (**Figure 4**). This indicates that the SVM framework utilizing importance and contribution measures to determine attribution and decision values was particularly well suited to this classification problem. Overall, the DescFits model presents a computationally light mathematical model to use for ALCA classification from routine EMR data·

The two cycles of feature selection (**Figure 2**) successfully removed highly correlated features (**Supplemental Figure S7**) to arrive at the final 4 features selected by the DescFits model. These 4 features continue to capture the clinical complexity of the 23-feature list including measures of renal function, cardiac stress, and hematologic state. Furthermore, 3 of the 4 selected features were parameters of fit to linear or exponential equations indicating that the temporal relationship of the data is important for determination of ALCA status. The intercepts selected by the DescFits 4 feature model are projected back to time 0. Replacing the intercept values with the raw values nearest time 0 (DescFits-Onco, blind AUC= 0·72) or diagnostic imaging (DescFits-Echo, blind AUC = 0·68) diminishes model performance on the blind set. By fitting patient data to model equations, the MISTIC SVM is able to capture the longitudinal relationship of data to a time aligned space.

When using AI and machine learning techniques for healthcare applications and decision making, explain-ability is key to adoption.^28^ The integrated gradient plots in **Figure 5 and Supplemental Figure S8** provide insight into how the decision value were reached by the DescFits 4 feature model. These figures illustrate how the DescFits model captures clinical relationships significant to the pathophysiology of cardiac amyloidosis. Test patients marked with stars in **Figure 5**, can be compared to patients with known ALCA status enabling use of MISTIC as an aid in clinical decision making. While there was a statistically significant (multiple unpaired t-test of the 4 selected features) difference between ALCA groups for chloride linear intercept (p = 0·0007), median BUN (p = 0·0070), and NT Pro BNP exponential intercept (p =0·01598) the role of serum electrophoresis beta linear intercept is unclear with traditional statistics. However, by plotting the integrated gradients against the feature values we can see that a relationship forms between serum electrophoresis beta linear intercept and the NT Pro BNP exponential intercept. The beta region of serum protein electrophoresis is a potential location of the M-spike in MM, yet its relationship to cardiac amyloidosis development has not been discussed in the literature. This could be a valuable area of future clinical study.

Existing models for classification of cardiac amyloidosis utilize additional data types such as ECG data ^20^ , echocardiography ^22^ , and proteomic data.^20,22,46^ Our model presents the simplicity of using only data which was routinely collected as part of a patient’s oncologic care. Further, the DescFits SVM model outperformed the models described by Lee (2023), Wu (2023), and Palstrom (2021). Agibetov et al 2020 created a gradient-boosted machine for classification of ALCA vs non-amyloidosis heart failure using routine laboratory data.^18^ However, this model relied upon imputation techniques for handling missing data and misses the shape of data over time. The missingness of EMR data often reflects a missing-not-at-random pattern, which violates the baseline assumptions for most imputation strategies.^24–27^

Fatigue is the most common symptom among ALCA patients ^7^ ; a symptom with a wide differential diagnosis, which may result in an increased time to ALCA diagnosis.^7^ Degree of cardiac involvement is the greatest predictor of survival^6,7^ with patients ultimately requiring heart transplant before end-stage renal disease develops.^6,7^ The current diagnostic algorithm for cardiac amyloidosis includes measuring NT-pro BNP, cardiac troponin-T, 24 urine proteins, and serum free light chains followed by biopsy positivity at another site (bone marrow or fat pad) and radiographic measures of concentric cardiomyopathy.^6,7,13^ However, interpretation of the cardiac markers used for prognostication can be obscured by declining renal function.^6,7^ Our model captures this complex cardio-renal clinical profile through both cardiac and renal longitudinal clinical features, which may increase sensitivity of detecting ALCA in MM patients. Implementing our model in the clinic could enable no-cost digital monitoring of MM patients throughout their treatment and decrease time to ALCA diagnosis.

Our model begins with the assumption that a patient has MM or MGUS and may thus be at risk for light chain ALCA. This model has not been tested on patients without these underlying assumptions and does not differentiate between ALCA and other forms of heart failure, which could be present in this patient population. Future work could include testing the DescFits model on patient data describing underlying hypertrophic cardiomyopathy unrelated to ALCA. Our current approach does not define the severity or stage of ALCA. However, future iterations of this work include building a regression model from the current DescFits SVM classifier to predict parameters of cardiac function including ejection fraction and left ventricular wall thickness. Testing the DescFits SVM model with a broader dataset is necessary to evaluate its performance at different clinical sites with different testing protocols. Fortunately, the DescFits model makes use of laboratory values which are commonly collected as part of complete blood counts and complete metabolic panels which may contribute to ease of site translation. Ultimately, this model could be further developed into an application or EMR plugin to assist with screening MM patients for ALCA in clinical settings. Developing an SVM regression model based on the DescFits model and MISTIC framework could provide a pretest computational screening for anatomic/physiologic cardiac measurements further reducing the healthcare burden of monitoring ALCA in MM patients. A regression model predicting cardiac function parameters such as ejection fraction or global longitudinal strain would complement the classification method by describing possible severity·

## ACKNOWLEDGEMENTS

This work was supported in part by the H. Lee Moffitt Cancer Center and Research Institute, an NCI-designated Comprehensive Cancer Center (P30-CA076292) Junior Scientist Partnership Award. Research reported in this publication was also supported by the UF Health Cancer Institute, supported in part by state appropriations provided in Fla. Stat. § 381·915 and the National Cancer Institute of the National Institutes of Health under Award Number P30CA247796. The content is solely the responsibility of the authors and does not necessarily represent the official views of the National Institutes of Health or the State of Florida. MAC is a trainee in the Medical Scientist Training Program supported by the National Institute of General Medical Sciences of the National Institutes of Health under award number T32GM163993.

## AUTHORS’ CONTRIBUTIONS

**MAC**: Methodology, Investigation, Formal Analysis, Visualization, Writing- original draft, review and editing

**HS**: Methodology, Software, Writing – review & editing

**MGF**: Data curation, Project administration, Writing – review & editing

**IBR**: Data curation, Funding acquisition, Project administration, Writing – review & editing

**CAK**: Conceptualization, Methodology, Software, Supervision, Project Administration, Funding Acquisition, Writing – review & editing

**MCF**: Conceptualization, Methodology, Investigation, Resources, Data Curation, Writing – Review & Editing, Supervision, Project Administration, Funding Acquisition

